# Changes in neural readout of response magnitude during auditory streaming do not correlate with behavioral choice in the auditory cortex

**DOI:** 10.1101/2022.06.14.496098

**Authors:** Taku Banno, Harry Shirley, Yonatan I. Fishman, Yale E. Cohen

**Affiliations:** Department of Otorhinolaryngology: Head and Neck Surgery, University of Pennsylvania School of Medicine, Philadelphia, PA 19104, USA; Departments of Neurology and Neuroscience, Albert Einstein College of Medicine, Bronx NY 10461, USA; Department of Neuroscience; Bioengineering, University of Pennsylvania, Philadelphia, PA 19104, USA

## Abstract

Although previous studies have identified neural mechanisms that may underlie auditory scene analysis, the relationship between these mechanisms and behavior remains elusive. To fill these gaps, we recorded multiunit activity (MUA) from the posterior and anterior auditory fields while monkeys participated in an auditory streaming task. We found that MUA magnitude was reduced as the streaming stimulus unfolded over time, and this reduction depended on the frequency difference between the tone bursts comprising the streaming stimulus. We then examined whether this frequency-dependent reduction in activity could be utilized by downstream neurons to read out “one stream” versus “two streams” and found that, as the frequency difference increased, an ideal observer consistently classified neural activity as “two streams”. However, because this classification was not modulated by the monkeys’ choices, it suggests that this activity may not reflect the segregation of stimuli into perceptually distinct auditory streams but may simply reflect bottom-up processes.

## Introduction

A fundamental goal of the auditory system is to transform auditory stimuli into perceptual representations of sound sources in the environment, a process commonly called “auditory scene analysis”^1^. An important component of this transformation is auditory-stream segregation. Stream segregation involves grouping stimuli that have similar spectrotemporal features (e.g., frequency) into one perceptual representation and the concomitant segregation of stimuli that have different features into different perceptual representations. For example, while listening to two interleaved temporal sequences of tone bursts, listeners often report hearing “one auditory stream” when the frequency difference between the tone bursts in the sequences is small. However, as the frequency difference between the tone bursts increases, listeners are more likely to report hearing “two auditory streams”. This segregation of multiple overlapping and interleaved sequences of tone bursts into distinct perceptual representations or auditory streams contributes to our ability to track a friend’s voice in a crowded restaurant or to follow the melody played by a particular instrument during a concert.

Although behavioral aspects of auditory-stream segregation have been studied extensively in humans^2–4^, the neural bases of auditory-stream segregation have yet to be fully elucidated. In part, this is due to the fact that neurophysiological studies of auditory streaming have been restricted to analyses of neural responses in the primary auditory cortex^5–11^ and its avian analogue^12^. Additionally, because most auditory streaming studies have been conducted in passively listening animals^5–8, 12^, the relationship between streaming behavior and simultaneous measures of neural activity still remains unclear.

To address these outstanding issues, we recorded neural activity in the posterior and anterior auditory fields of the auditory cortex, which include both the primary and non-primary auditory cortices, while rhesus monkeys performed an auditory streaming task^13^. In this streaming task, monkeys listened to a temporal sequence of interleaved low- and high-frequency tone bursts and detected a deviantly loud “target” tone burst that was embedded in the low-frequency sequence. Because this deviant target could only be reliably detected when the low-frequency sequence was segregated from the high-frequency sequence, the task provided an objective measure of auditory streaming^13^.

Consistent with previous reports^7, 14^, we found that as the streaming stimulus unfolded over time, the magnitude of MUA was reduced, and this reduction depended on the frequency difference between the low- and high-frequency tone bursts of the streaming stimulus. We then asked whether this frequency-dependent reduction in activity could be used by downstream neurons to read out “one stream” versus “two streams” and found that as frequency difference between the low- and high-frequency tone bursts increased, an ideal observer consistently categorized neural activity as “two stream”, consistent with human reports. However, because this classification was not modulated by the monkeys’ choices, it suggests that changes in the magnitude of neural activity, at least in the brain regions that we tested, does not reflect perceptual stream segregation but may simply reflect bottom-up processes.

## Materials and Methods

We obtained multilaminar extracellular recordings in the auditory cortex of two adult male macaque monkeys (*Macaca mulatta*; monkey D and monkey C) while they performed a behavioral task that provided an objective measure of auditory streaming. The Institutional Animal Care and Use Committee of the University of Pennsylvania reviewed and approved all the procedures and protocols. We conducted all surgeries under general anesthesia with aseptic techniques.

### Chamber placement and Identification of the posterior and anterior auditory fields

We identified the location of the auditory cortex by structural MRI scans of each monkey’s brain and then used these images to stereotaxically implant a recording chamber over the dorsal surface of the monkey’s skull^15, 16^. We oriented the recording chamber so that a multi-channel electrode penetrated the auditory cortex roughly perpendicular to the cortical lamina to enable sampling of neural activity across the entire laminar extent of auditory cortex^17, 18^.

We functionally defined different auditory fields according to their tonotopic organization. To do this, in each session, we generated each neuron’s spectrotemporal receptive field (STRF; details below) and used the STRF to estimate the neuron’s “best frequency” (BF); the BF was defined as the frequency value that elicited the highest neural firing rate (see below for details). We constructed tonotopic maps for each monkey by plotting BF as a function of the anterior-posterior position within the recording chamber. As in a previous report^19^, we defined the border between posterior and anterior auditory fields as the location of the low-frequency gradient reversal. The posterior auditory field mainly overlaps with the primary auditory cortex (A1) but may also include recording sites in ML, MM, CL, and CM. The anterior auditory field mainly overlaps with rostral core and belt auditory areas including R, RM, and AL^20, 21^.

### Experimental chamber

We conducted behavioral and recording sessions in a darkened room with echo- and sound-attenuating walls. We seated a monkey in a primate chair with a non-invasive head restraint^22^. During the auditory-streaming task, the monkeys indicated their behavioral report by releasing a touch-sensitive lever (monkey D) or moving a joystick (monkey C). We generated (Matlab, The Mathworks) and delivered (RZ8, TDT Inc or e22, Lynx Inc.) calibrated auditory stimuli via a free-field speaker (TR-3 [Anthony Gallo Acoustics] or MSP5 [Yamaha]). We positioned the speaker at each monkey’s eye level, which was ∼1 m in front of them.

### Auditory-streaming task

The auditory-streaming task is a modified version of a task used to evaluate auditory-stream segregation in human listeners^13^. In our task, monkeys listened to two interleaved sequences of tone bursts (50-ms duration; 25-ms inter-burst interval). One sequence consisted of “low frequency” (L) tone bursts, and the other sequence consisted of “high frequency” (H) tone bursts. The frequency value of the low-frequency tone-burst sequence was held constant in a session, whereas we varied trial-by-trial the frequency value of the high-frequency tone-burst sequence. The frequency range of the high-frequency tone bursts was 1-24 semitones above the low-frequency value (Fig. 1a). We presented these tone bursts as a repeating sequence of “low-high-high” (L-H-H) triplets (Fig. 1a), analogous to “ABB” triplets used in a previous streaming study ^13^.

**Figure 1:**
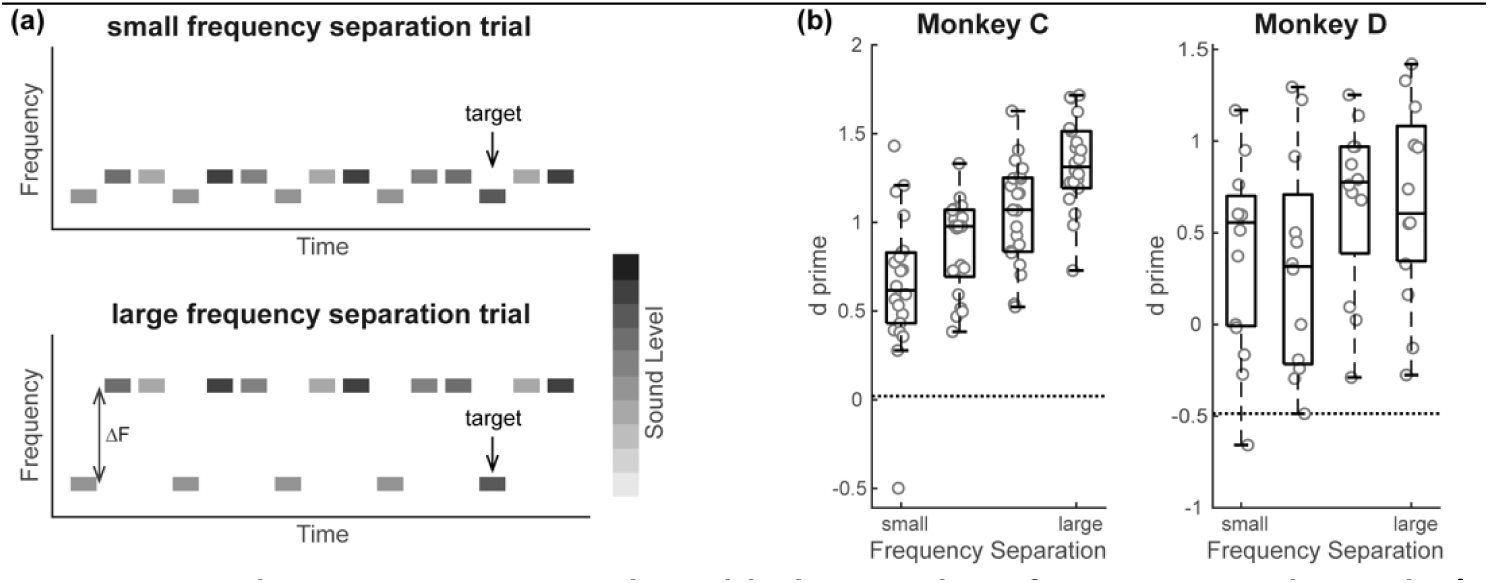
Auditory streaming task and behavioral performance to the task. **(a)** Schematic illustration of the auditory streaming task. Each rectangle indicates low-(L) and high-(H) frequency tone burst presented as repeating L-H-H triplets. The rectangle’s shading represents the relative sound level of the tone bursts: the darker the color, the louder the sound level (see legend). The task’s goal was for a monkey to detect a relatively louder “deviant” target in the low-frequency tone burst sequence (as indicated in the schematic). Task difficulty was titrated by varying the frequency difference (ΔF) between the low- and high-frequency tone burst bursts. **(b)** d’ as a function of ΔF for monkey C and D, respectively. The gray circles indicate the d’ from each recording session, and the box plot indicates the median and upper and lower quantiles of the d’ across sessions. The dotted line indicates the mean of the bootstrapped null distribution for each monkey.

We presented the low-frequency tone bursts at two sound levels: 52 or 68 dB SPL. In each trial, only one of the low-frequency tone bursts was presented at 68 dB, whereas all the other low-frequency tone bursts in the trial were presented at 52 dB. This “deviantly” loud 68-dB tone burst was the target stimulus. The monkeys reported its presence by releasing a lever or moving a joystick. The high-frequency tone bursts were presented at one of four randomly chosen sound levels, which spanned a range extending above and below the levels of the low-frequency bursts: 47, 57, 62, and 72 dB SPL (Fig. 1a).

We titrated task difficulty by varying the frequency difference (ΔF) between the low- and high-frequency tone-burst sequences. Because previous work suggests that monkeys report streaming stimuli in a manner comparable to human listeners^23–25^, we hypothesized that changes in ΔF would also affect the monkeys’ performance in a similar way as human listeners^13^. Specifically, when the ΔF between the two sequences was small, the low- and high-frequency tone-burst sequences would be perceptually integrated into a single auditory stream. As a result, the sound level of the target would be within the variability of the sound levels of the low- and high-frequency tone bursts and would be difficult to detect as being “deviantly loud”. In contrast, when the ΔF between the sequences was large, the sequences would perceptually segregate into two auditory streams and the target would be more readily detectable: that is, the louder sound level of the 68-dB target would be more salient relative to the softer 52-dB tone bursts. The smallest and largest ΔF values varied across recording sessions (typically 1-24 semitones for monkey D and 2-24 semitones for monkey C).

To minimize the possibility that monkeys could anticipate target onset, we randomized the time of target onset between 675 and 2025 ms, relative to sequence onset. In other words, the target tone burst could appear in any position between the 4^th^ and the 10^th^ L-H-H triplet. The maximum number of triplets in a sequence was 13.

When the monkey responded within a specified temporal window following target onset (monkey D: window = 650 ms; median response time between target onset and movement: 431 ± 81 ms; monkey C: window = 800 ms; median response time: 517 ± 110 ms; differences in the temporal windows were based on the individual monkey’s idiosyncratic behavior with their respective manipulandum), we considered the trial to be a “hit”. If the monkey responded after this response window or did not respond at all, we considered the trial to be a “miss”. A “false alarm” occurred when the monkey responded before target onset. Conservatively, we also considered very rapid responses (<200 ms for monkey D and <250 ms for monkey C) as false alarms. We rewarded monkeys only on hit trials. For both miss and false-alarm trials, we added a timeout (1500-2500 ms) to the inter-trial interval following these incorrect trials.

### Neurophysiological recording strategy

We inserted a linear multi-channel electrode (16-channel v-probe, 150-µm spacing between channels; or a 24-channel s-probe, 100-µm spacing between channels; Plexon Inc.) into the brain through a stainless-steel guide tube. The electrode was then advanced gradually with a microdrive (NAN Instruments). While advancing the electrode, we presented an auditory “search” stimulus (100-ms Gaussian noise burst; 10-ms cos^2^ rise and fall times; 900 ms inter-burst-interval; 50-kHz sampling rate) to identify auditory responsive sites. Neural signals were amplified (PZ2 and PZ5, TDT Inc.), digitized, and stored sampling rate: 24.4 kHz; RZ2, TDT Inc.) for online and offline analyses.

Guided by laminar profiles of multiunit activity (MUA) and current-source-density (CSD) that were evoked by the search stimulus, we finely adjusted the electrode’s depth so that the largest stimulus-evoked MUA, which typically coincided with a prominent initial current sink in the CSD profile, was positioned in the electrode’s middle channels^9, 18^. To minimize electrode drift during a recording session and artifactual components in the CSD profile (e.g., due to mechanical compression or distortion of the cortical tissue by the electrode), we retracted the electrode by ∼200-450 µm and allowed the tissue to stabilize for ≥30 minutes before further data collection. After we finished collecting task-related data, we obtained another set of MUA and CSD profiles to confirm electrode stability over the recording session.

During this ∼30-minute stabilization period, we characterized the spectrotemporal receptive field (STRF) of the recorded neurons by passively presenting a dynamic moving ripple (DMR) auditory stimulus ^26, 27^. The DMR is a continuous time-varying broadband noise stimulus that covers the frequency range between 0.1 and 35 kHz (5-min duration; 65 dB spectrum level per ⅓ octave; 96-kHz sampling rate; 24-bit resolution). At any instant of time, the stimulus had a sinusoidal spectrum; the spectral modulation frequency (0-4 cycles/octave) determined the density of the spectral peaks. The peak-to-peak amplitude of the ripple was 30 dB. The temporal modulation frequency (0-50 Hz) controlled the stimulus’ temporal modulations. Spectral and temporal parameters were varied randomly, dynamically, and independently; the maximum rates of change for these spectral and temporal parameters were 0.25 Hz and 1.2 Hz, respectively. We derived the STRF by averaging the spectrotemporal envelope of the DMR relative to the time of each spike recorded at each electrode channel^26, 27^. We considered the frequency value corresponding to the STRF peak as the BF of the electrode channel.

The BF defined the frequency values of the tone-burst sequences presented in the auditory-streaming task. When the BF was <3.5 kHz, we set the value of the low-frequency tone-burst sequence to the BF value. When the BF was >3.5 kHz, we set one of the high-frequency values (usually at the largest frequency difference [ΔF]) to this BF value; we do not report neural data from these higher-BF sites. The monkey then performed the auditory-streaming task. On a trial-by-trial basis, we randomly varied the ΔF and the target onset time.

### Behavioral data analysis

In each recording session, we calculated the hit, miss, and false-alarm rates. Behavioral d’ was defined as the difference between the z-transform of the hit and false-alarm rates. We calculated d’ as a function of the frequency difference between the tone-burst sequences. Additionally, to estimate the false-positive rate of the d’ values, we used a bootstrap randomization procedure to generate null distributions.

### Neural data analysis

#### Extraction of MUA envelope and identification of stimulus-evoked MUA

Our neural-data analyses focused on the time-varying MUA to facilitate comparison between our findings and those of previous streaming studies^5, 6, 28^. MUA and single-unit techniques have been shown to yield similar response properties, whereas MUA is generally more stable than single-unit activity^29–31^.

For each trial and for each electrode channel, we extracted the MUA envelope by first bandpass filtering the neural signal (0.5-3.0 kHz), full-wave rectifying it, and finally low-pass filtering (0.6-kHz cutoff frequency) this signal^5, 18, 30, 32, 33^. We then obtained an integrated MUA response to the tone bursts by summing, trial-by-trial, the MUA envelope over a 75-ms window that included the 50-ms duration of each tone burst and the 25-ms silent gap that followed the offset of each tone burst. This summed MUA was then z-scored based on a distribution of “baseline” values generated from a random sampling of neural activity from 75-ms windows comprising the -1000 ms to -500 ms period that preceded the onset of the tone-burst sequence. We sampled these baseline values across all trial conditions so that the conditional differences in baseline activity would not affect the z-score normalization.

We only report data from electrode channels that showed “stimulus-evoked” MUA. To determine whether a channel showed stimulus-evoked MUA, we averaged the trial-by-trial z-scored MUA (zMUA) as a function of the frequency difference between the low-and high-frequency tone bursts. If at least one of the zMUA values from the first L-H-H triplet was >1.96 (95% confidence level of z-score value), we considered the zMUA for the channel as “stimulus evoked”.

#### Receiver-operating-characteristic (ROC) analyses of auditory streaming

We used ROC analyses to evaluate the degree to which an ideal observer could discriminate between two distributions of trial-by-trial zMUA values. The area under the ROC curve (auROC) is the probability that an ideal observer could discriminate between the two distributions^34^. In one analysis, we analyzed the differences in zMUA amplitude elicited by the low versus high-frequency tone bursts as the streaming sequence unfolded over time. To do this, we generated one distribution of single-trial zMUA values in response to the low-frequency tone burst and a second distribution of zMUA values in response to the first high-frequency tone burst in each L-H-H triplet. An auROC value of 0.5 indicates that an ideal observer could not discriminate between those two distributions. Values different from 0.5 indicate that an ideal observer could discriminate between these distributions with values greater than 0.5 indicating that mean value of the low-frequency zMUA distribution was larger than the mean value of the high-frequency zMUA distribution. We conducted this analysis as a function of L-H-H triplet position (in which the “first” triplet was the first one presented in a trial, etc.) and ΔF.

#### Statistical analyses of CI, FSI and BMI

We used non-parametric statistics and post-hoc comparisons to evaluate null hypotheses. In all statistical tests, we rejected the null hypothesis at p< 0.05. The false-discovery rate was corrected by the Benjamini– Hochberg procedure^35^.

### Data and code availability

Raw data from Figures were deposited on Mendeley at doi:10.17632/rth4ssfc4n.1

## Results

### The auditory-streaming task: behavioral data

The auditory-streaming task consisted of temporally interleaved sequences of low-(L) and high-frequency (H) tone bursts that were presented as an “L-H-H” triplet (Fig. 1a). On each trial, all the low-frequency tone bursts had the same sound level except for one deviantly loud “target” tone burst. The sound levels of each high-frequency tone burst were pulled randomly from a distribution of values that were louder and softer than those of the low-frequency tone bursts. Based on a human psychophysical study^13^, we reasoned that when the frequency difference (ΔF) between the low- and high-frequency was small (large), target detection would be harder (easier). A monkey used a manipulandum to report the target tone burst and receive a reward. See **Materials and Methods** for more information on the stimulus and task design.

Both monkeys’ behavioral sensitivity (d’) was significantly greater than their bootstrapped chance level (95% confidence interval of bootstrapped null distribution. Monkey D: 0.016 – 0.022; monkey D: - 0.49 – -0.47), especially in the larger ΔF conditions (Fig. 1b and 1c). Furthermore, monkey C’s performance significantly improved with ΔF (Fig. 1b, p=8.40×10^-8^; H_0_: d’ was the same across all ΔFs; Kruskal-Wallis test).

For monkey D, we could not identify a reliable relationship between d’ and ΔF (Fig. 1c, p=0.14; H_0_: d’ was the same across all ΔFs; Kruskal-Wallis test). On average, these behavioral results are consistent with the hypothesis that, like human listeners^13^, target detection is more reliable in the monkeys when the low-frequency tone-burst sequence is perceptually segregated, which occurs at large ΔFs, from the high-frequency tone-burst sequence.

### Posterior and anterior auditory fields are defined by tonotopic gradients

We mapped each site’s best frequency (BF) (see Fig. 2a for example STRFs and their corresponding BFs) along the anterior-posterior axis of the recording chamber and found a reversal in the tonotopic gradient in both monkeys (Fig. 2b and 2c). Accordingly, we separated our recording sites into “posterior” and “anterior” auditory fields by setting a border at the low-frequency tonotopic reversal (Fig. 2b and 2c)^19^. The posterior auditory field includes primary auditory cortex (A1) and may also include the posterior belt auditory cortex (CL, CM, MM and ML), whereas the anterior auditory field includes areas in the rostral core (R) and the anterior belt auditory cortices (RM and AL)^20, 21^. We could not identify any other reliable neurophysiological markers that allowed us to further resolve our recording sites (such as mediolateral position) into specific areas of the core or belt auditory cortex.

**Figure 2:**
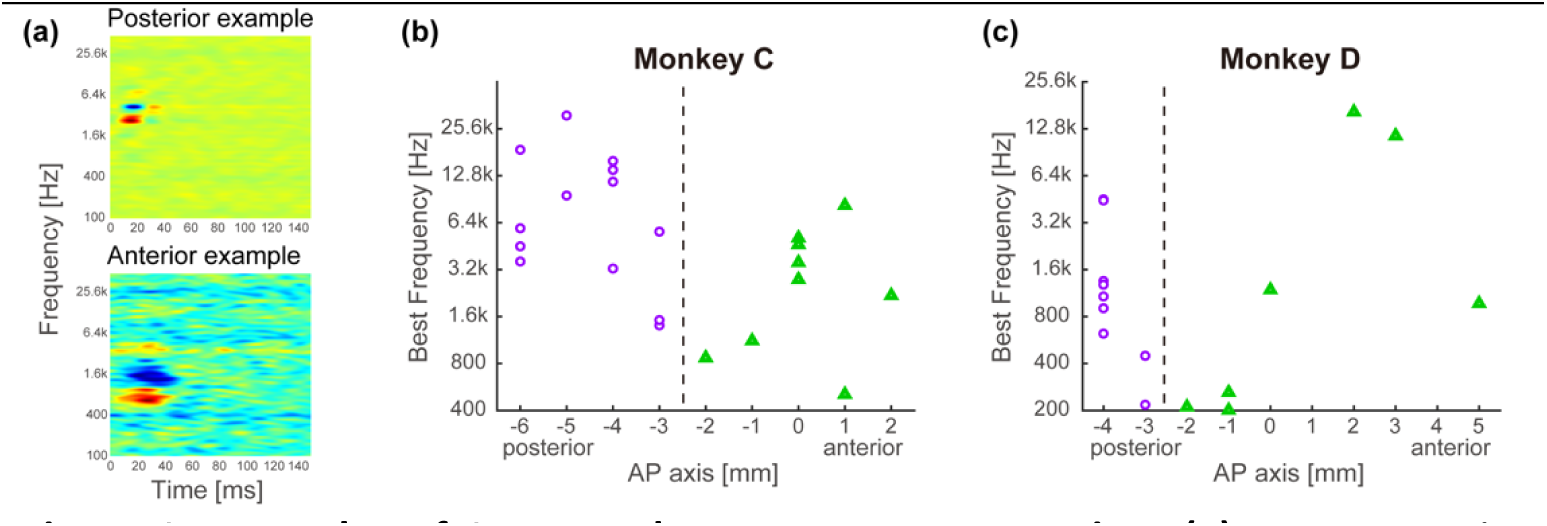
Examples of STRFs and tonotopy reconstruction. **(a)** Representative spectrotemporal receptive fields (STRFs) obtained from the **(top)** posterior and **bottom)** anterior auditory fields. The x-axis is aligned relative to stimulus onset. Hotter/cooler colors indicate increased/decreased firing rates, respectively. **(b-c)** Reconstructions of tonotopy in the posterior (purple circles) and anterior (green triangles) auditory fields in monkeys C and D. The best frequency of neural activity i.e., the frequency value that elicited the highest neural firing rate from the STRFs) increased at more posterior positions in the posterior auditory field, whereas those in the anterior auditory field increased at more anterior positions. The border between the posterior and anterior auditory fields was inferred from the location of the frequency-gradient reversal (dotted line).

### MUA responses were modulated by time and ΔF

We report findings based on stimulus-evoked neural activity recorded from 147 electrode channels in 18 electrode penetrations (see **Materials and Methods**). 77 of these recordings came from the posterior auditory field, and 70 came from the anterior auditory field.

Figure 3 shows an example of MUA recorded during the auditory-streaming task. Three main findings emerge from this sample recording. First, as expected, because the low-frequency tone bursts were set to the site’s BF, they elicited robust MUA responses. In contrast, MUA elicited by the high-frequency tone bursts decreased in amplitude as the frequency of the high-frequency tone bursts moved away from the BF (i.e., at larger ΔFs). Second, MUA decreased in amplitude as the stimulus sequence unfolded over time.

**Figure 3:**
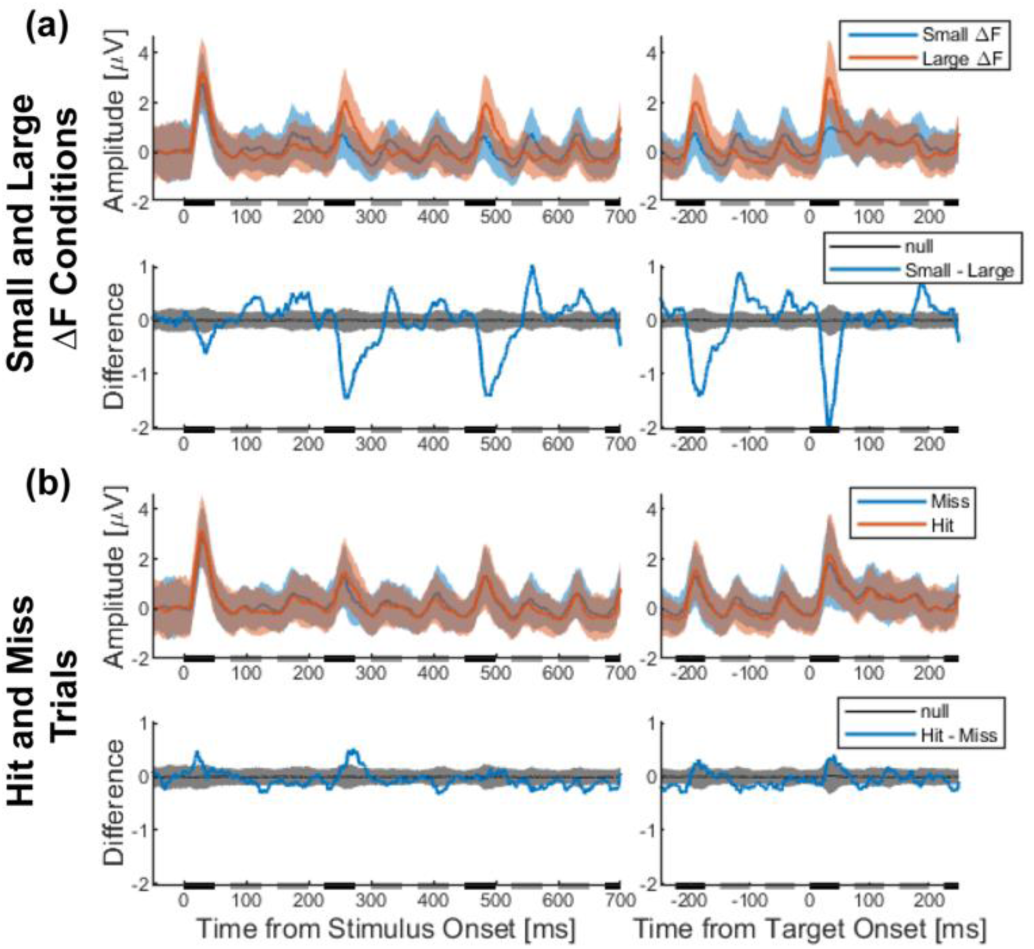
Example of MUA during the auditory-streaming task. **(a, top)** A MUA response profile from the posterior auditory field. The profile shows the mean MUA from trials with a 1-semitone difference and an 8-semitone difference. The rectangles on the horizontal axis indicate the presentation times of low-(black) and high-(gray) frequency tone bursts. **(a, bottom)** The difference in mean MUA between different stimulus (small ΔF versus large ΔF; blue curve). The bootstrapped null distribution (mean ± 95% confidence intervals) is plotted in grey. **(b)** The data from the same recording site but reorganized as a function choice (hit versus miss). The organization of the curves in both panels is the same as in **(a).**

In this figure, this was most noticeable for the responses to the low-frequency tone bursts. Finally, MUA was comparable on hit and miss trials and thus did not appear to be substantially modulated by the monkey’s behavior (Fig. 3b). Below, we quantify and expand on all three of these observations.

### MUA decreases in amplitude over time and as a function of ΔF

Micheyl and colleagues suggested a model of auditory-stream perception that incorporates some form of neural habituation or adaptation, whereby neural responses decrease in amplitude over time^7, 36^. To test this proposal in our data, we first quantified whether neural representations of the low- and high-frequency tone bursts decrease in amplitude over time, independent of the monkeys’ behavioral choice (Fig. 4). In general, in the posterior auditory field, MUA response amplitude decreased over time (i.e., triplet position; see Fig. 1a) in response to the low-frequency tone burst (p = 4.59 x 10^-18^, H_0_: zMUA was the same at all triplet positions; Scheirer-Ray-Hare test) and for the first high-frequency tone burst (p = 3.16 x 10^-7^, H_0_: zMUA was the same at all triplet positions; Scheirer-Ray-Hare test) but not in response to the second high-frequency tone burst (p = 0.28, H_0_: zMUA was the same at all triplet positions; Scheirer-Ray-Hare test). We also found a main effect of ΔF on zMUA in response to each of the tone bursts in a triplet (p<0.05, H_0_: zMUA was the same across all ΔFs; Scheirer-Ray-Hare test).

**Figure 4:**
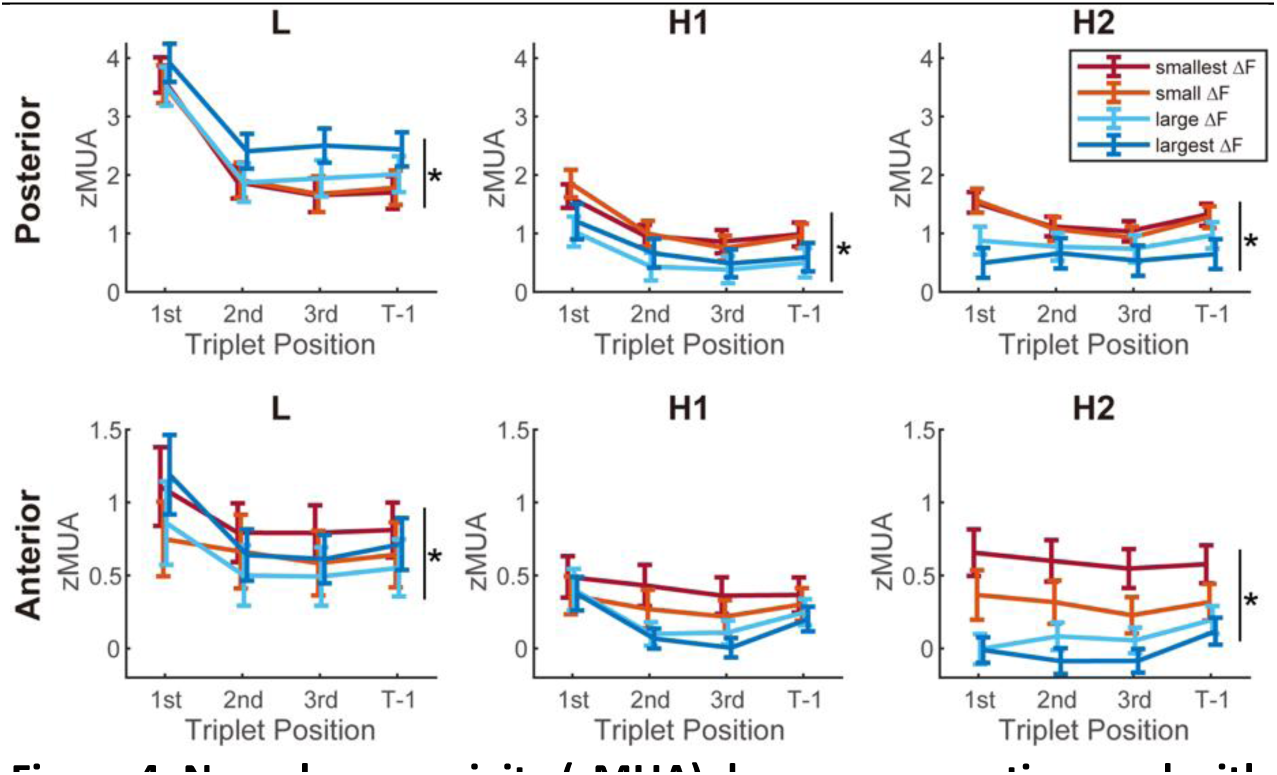
Neural responsivity (zMUA) decreases over time and with ΔF. zMUA values are shown as a function of time and ΔF (see legend); data are aligned relative to the first three (“1st”, “2nd”, and “3rd”) triplet positions and for the triplet immediately preceding the target (i.e., “T-1”). zMUA values in the **top row** were calculated from data collected in the posterior auditory field, whereas those in the **bottom row** were calculated from data collected in anterior auditory field. Data in the **first column** are zMUA values in response to the low-frequency tone burst (L) of the L-H-H triplet; those in the **middle column** are in response to the first high-frequency tone burst (H1); whereas those in the **last column** are in response to the second high-frequency tone burst (H2). zMUA values are reported across trials, regardless of behavioral choice. Data points indicate the population mean and standard error. The asterisks indicate when the null hypothesis was rejected (p<0.05, H0: zMUA was the same across all ΔFs; Scheirer-Ray-Hare test).

*MUA decreases in amplitude over time and as a function of ΔF* Micheyl and colleagues suggested a model of auditory-stream perception that incorporates some form of neural habituation or adaptation, whereby neural responses decrease in amplitude over time^7, 36^. To test this proposal in our data, we first quantified whether neural representations of the low- and high-frequency tone bursts decrease in amplitude over time, independent of the monkeys’ behavioral choice (Fig. 4). In general, in the posterior auditory field, MUA response amplitude decreased over time (i.e., triplet position; see Fig. 1a) in response to the low-frequency tone burst (p = 4.59 x 10^-18^, H_0_: zMUA was the same at all triplet positions; Scheirer-Ray-Hare test) and for the first high-frequency tone burst (p = 3.16 x 10^-7^, H_0_: zMUA was the same at all triplet positions; Scheirer-Ray-Hare test) but not in response to the second high-frequency tone burst (p = 0.28, H_0_: zMUA was the same at all triplet positions; Scheirer-Ray-Hare test). We also found a main effect of ΔF on zMUA in response to each of the tone bursts in a triplet (p<0.05, H_0_: zMUA was the same across all ΔFs; Scheirer-Ray-Hare test).

In contrast, in the anterior auditory field, the effect of triplet position on the zMUA response amplitude was weaker and was only significant for the first high-frequency tone burst (low-frequency tone burst: p = 0.70, first high-frequency tone burst: p = 0.02, and second high-frequency tone burst: p = 0.33; H_0_: zMUA was the same at all triplet positions; Scheirer-Ray-Hare test). We also found that ΔF modulated zMUA in response to the low-frequency (p = 0.002; H_0_: zMUA was the same across all ΔFs; Scheirer-Ray-Hare test) and second high-frequency (p = 2.75 x 10^-13^; Scheirer-Ray-Hare test) tone bursts but not the first high-frequency tone burst (p = 0.07; Scheirer-Ray-Hare test).

To further quantify these observations, we used a receiver-operating characteristic (ROC) analysis (see **Materials and Methods**) to calculate the degree to which an ideal observer could differentiate between zMUA that was elicited by the low-frequency tone burst and the first high-frequency tone burst in an L-H-H triplet. Initially, in the posterior auditory field (Fig. 5a), across all ΔFs, auROC values were significantly greater than 0.5 indicating that the low-frequency (BF) tone bursts elicited more activity (on average) than the first high-frequency tone burst (Table 2). This initial auROC value is consistent with the frequency tuning properties of auditory-cortex neurons, which respond more to tone bursts with frequencies at or near their BF (the low-frequency tone bursts) than to tone bursts with frequencies away from the BF (the high-frequency tone bursts). As the tone-burst sequence unfolded over time, the auROC values from smaller ΔF trials decreased significantly over time (Table 1; p<0.05, H_0_: auROC values were the same at all triplet positions; Kruskal-Wallis test) but still remained significantly greater than 0.5 (Table 2; p<0.05, H_0_: auROC values were the same as 0.5; Wilcoxon signed-rank test with FDR correction). In contrast, in larger ΔF trials, auROC values remained constant across time (Table 1; p=0.11 and p=0.24 for larger ΔF trials, H_0_: auROC values were the the same at all triplet positions; Kruskal-Wallis test). A somewhat different finding was seen in the anterior auditory field (Fig. 5b). Here, auROC values in both small and large ΔF trials–except in the smallest ΔF trials, decreased over time (Table 1). The auROC values on large ΔF trials were still larger than on small ΔF trials. Overall, our data suggest that in both auditory fields, the difference between zMUA, which was elicited by the low- and high-frequency tone bursts, generally decreases as the streaming sequences unfold over time, as compared with the discriminability observed for responses evoked by the initial triplet in the sequences.

**Figure 5:**
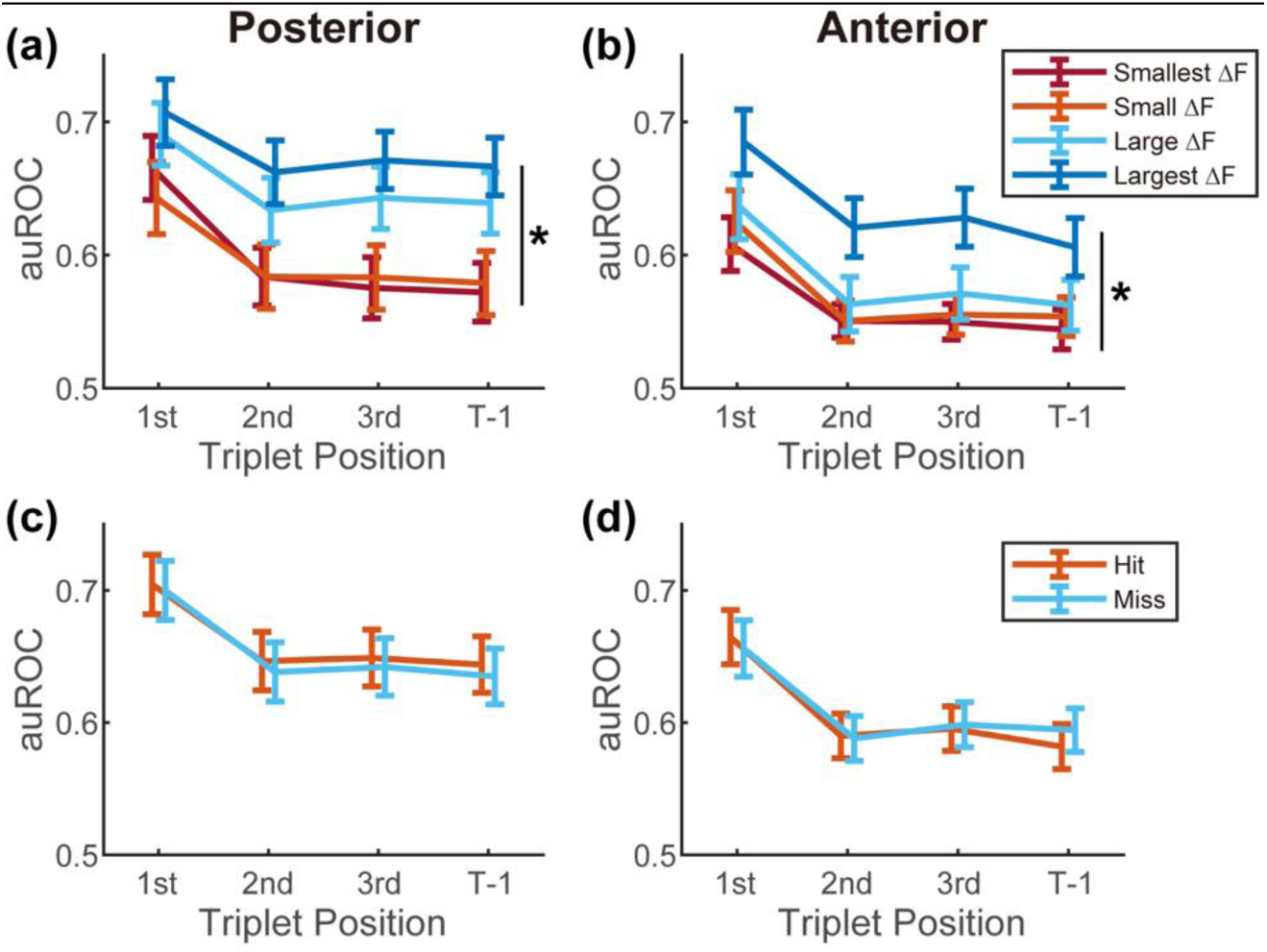
Sensitivity of neural responsivity to the low- and first high-frequency tone bursts. auROC values, which were generated from the single-trial zMUA distributions in response to the low-frequency tone burst and the first high-frequency tone burst of the L-H-H triplet, are shown as a function of time; data are aligned relative to the first three (“1^st^”, “2^nd^”, and “3^rd^”) triplet positions and for the triplet immediately preceding the target (i.e., “T-1”). auROC values from data collected in the posterior auditory field are shown in **(a)**, whereas auROC values from the anterior auditory field are shown in **(b)**. These data are plotted as a function of ΔF (see legend) but independent of behavioral choice. Data in **(c)** and **(d)** show auROC values from data generated on hit and miss trials, respectively, across all ΔFs. In each panel, data points indicate the population mean and standard error. The asterisks indicate when the null hypothesis was rejected (p<0.05, H_0_: auROC values are the same across ΔF or behavioral choice; Scheirer-Ray-Hare test).

**Table 1;.**
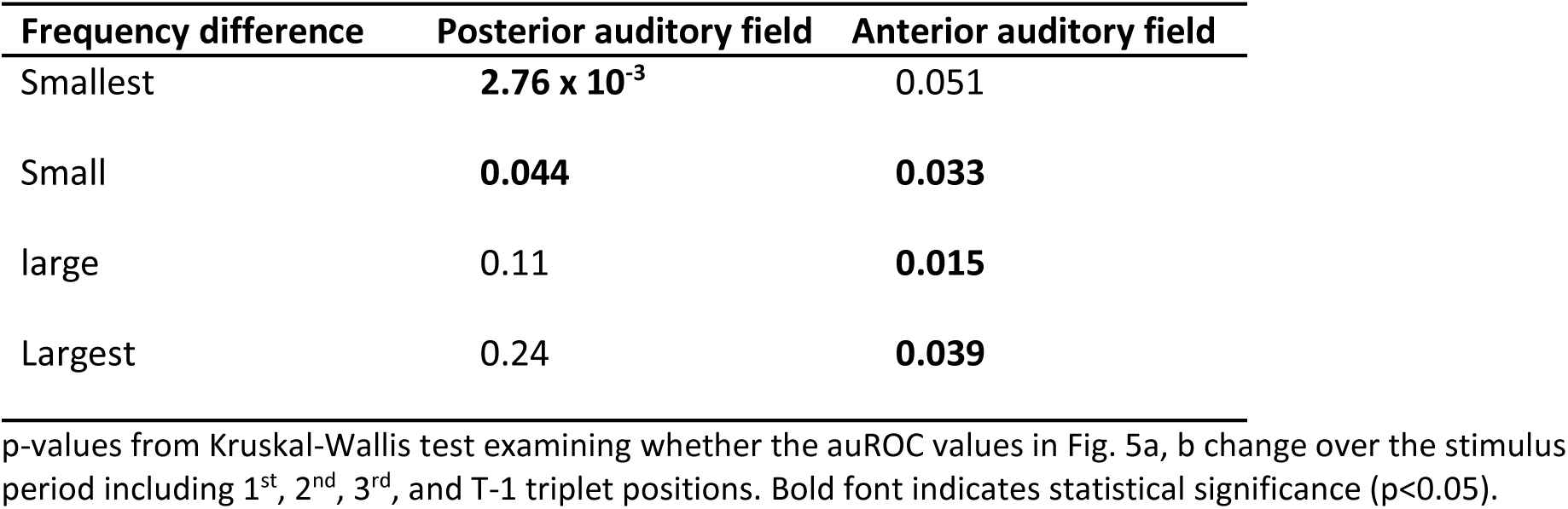
Statistical test examining the time course of stimulus-related zMUA modulation

| Frequency difference | Posterior auditory field | Anterior auditory field |
| --- | --- | --- |
| Smallest | **2.76 x 10<sup>-3</sup>** | 0.051 |
| Small | **0.044** | **0.033** |
| large | 0.11 | **0.015** |
| Largest | 0.24 | **0.039** |

p-values from Kruskal-Wallis test examining whether the auROC values in Fig. 5a, b change over the stimulus period including 1<sup>st</sup>, 2<sup>nd</sup>, 3<sup>rd</sup>, and T-1 triplet positions. Bold font indicates statistical significance ( $p < 0.05$ ).

**Table 2;.**
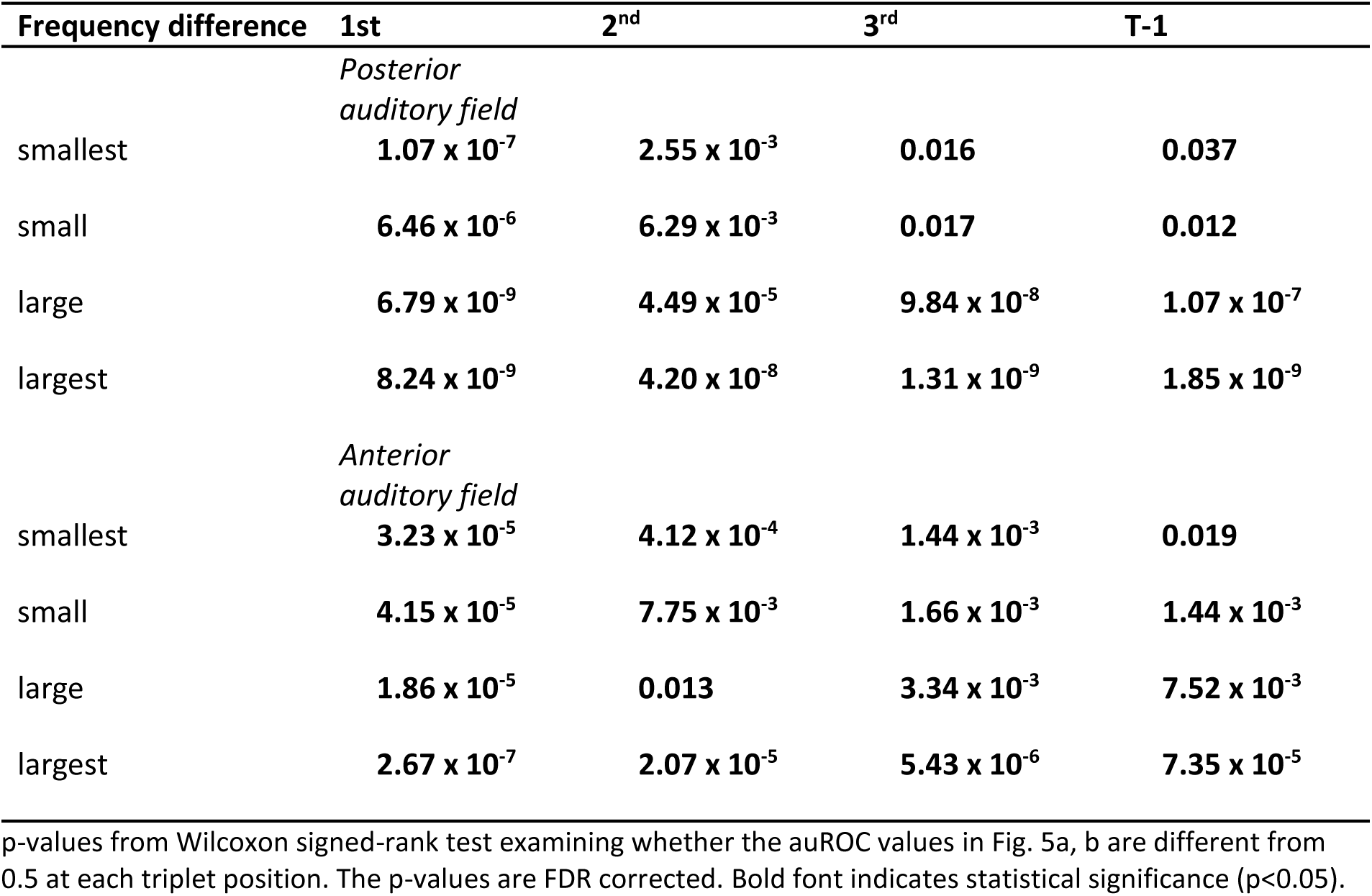
Statistical test examining zMUA stimulus sensitivity at each triplet position

When we collapsed across ΔF but analyzed neural activity as a function of choice (hit versus miss trials), we found that the ability of an ideal observer to discriminate between zMUA elicited on low- and high-frequency trials decreased over time (p = 2.10 x 10^-3^ in posterior, p = 3.55 x 10^-4^ in anterior; H_0_: auROC values were the same at all triplet positions; Scheirer-Ray-Hare test) but was not dependent on the monkeys’ choices (p = 0.43 in the posterior auditory field, p = 0.98 in the anterior auditory field; H_0_: auROC values were the same for hit and miss trials; Scheirer-Ray-Hare test) (Fig. 5c, d).

### A potential mechanism to read-out “one stream” versus “two streams”

Next, we considered whether a time- and frequency-dependent reduction in MUA response amplitude might enable downstream neurons to decide whether the tone-burst sequences could be interpreted as “one stream” or “two streams” depending on whether the low- and high-frequency tone-burst responses both exceed a threshold value^7^. Reports of “one stream” would be more likely to occur on small-ΔF trials when both the low- and high-frequency tone-bursts are close to the BF of the recorded neurons, and hence both elicit comparatively large responses. In contrast, a listener would report “two streams” when only the response to the low-frequency tone burst would exceed this threshold, which would be more likely to occur on large-ΔF trials when the high-frequency tone burst is far from the BF and hence elicits smaller responses.

For this analysis, we first found the zMUA value that optimally separated the distributions of zMUA values elicited by the low- and high-frequency tone bursts (i.e., the decision boundary); initially, we included both hit and miss trials. Both distributions included data generated in response to the first triplet through the one immediately preceding the target triplet and across all ΔFs in each recording session. (Once this boundary was determined, it was fixed and was not recalculated for different time points nor for different ΔFs.) Next, as a function of time and ΔF, we evaluated whether the mean values of the distributions of low-frequency and high-frequency zMUA responses were above or below this threshold. If both were above the threshold, the responses were classified as representing “one stream”. On the other hand, if only the mean of the low-frequency distribution was above the threshold, the responses were classified as representing “two streams”. Across trials, ΔF, and sessions, we calculated the probability of “two streams”.

In the posterior auditory field, we found that the probability that the ideal-observer model classified neural activity as “two streams” reports increased significantly as ΔF increased, (Fig. 6a; p = 4.40 x 10^-7^, H_0_: probability of “two stream” reports is same across all ΔFs; Scheirer-Ray-Hare test). However, triplet position did not significantly affect the probability of the neural model classifying neural activity as “two streams” (p = 0.06, H_0_: probability of “two stream” reports is same at all triplet positions; Scheirer-Ray-Hare test). A similar set of findings was observed in the anterior auditory field (Fig. 6b; p = 4.00 x 10^-21^, p = 0.07, H_0_: probability of “two stream” reports is same across ΔF and is the same at all triplet positions, respectively; Scheirer-Ray-Hare test). Further, when we analyzed hit and miss trials separately (Fig. 6c, 6d), we found that the monkeys’ behavioral choices had minimal impact on the neural responses leading to a “one stream” versus “two streams” categorization (p = 0.38 for the posterior auditory field, p = 0.54 for the anterior auditory field; H_0_: probability of “two stream” reports is same for behavioral choice; Scheirer-Ray-Hare test).

**Figure 6:**
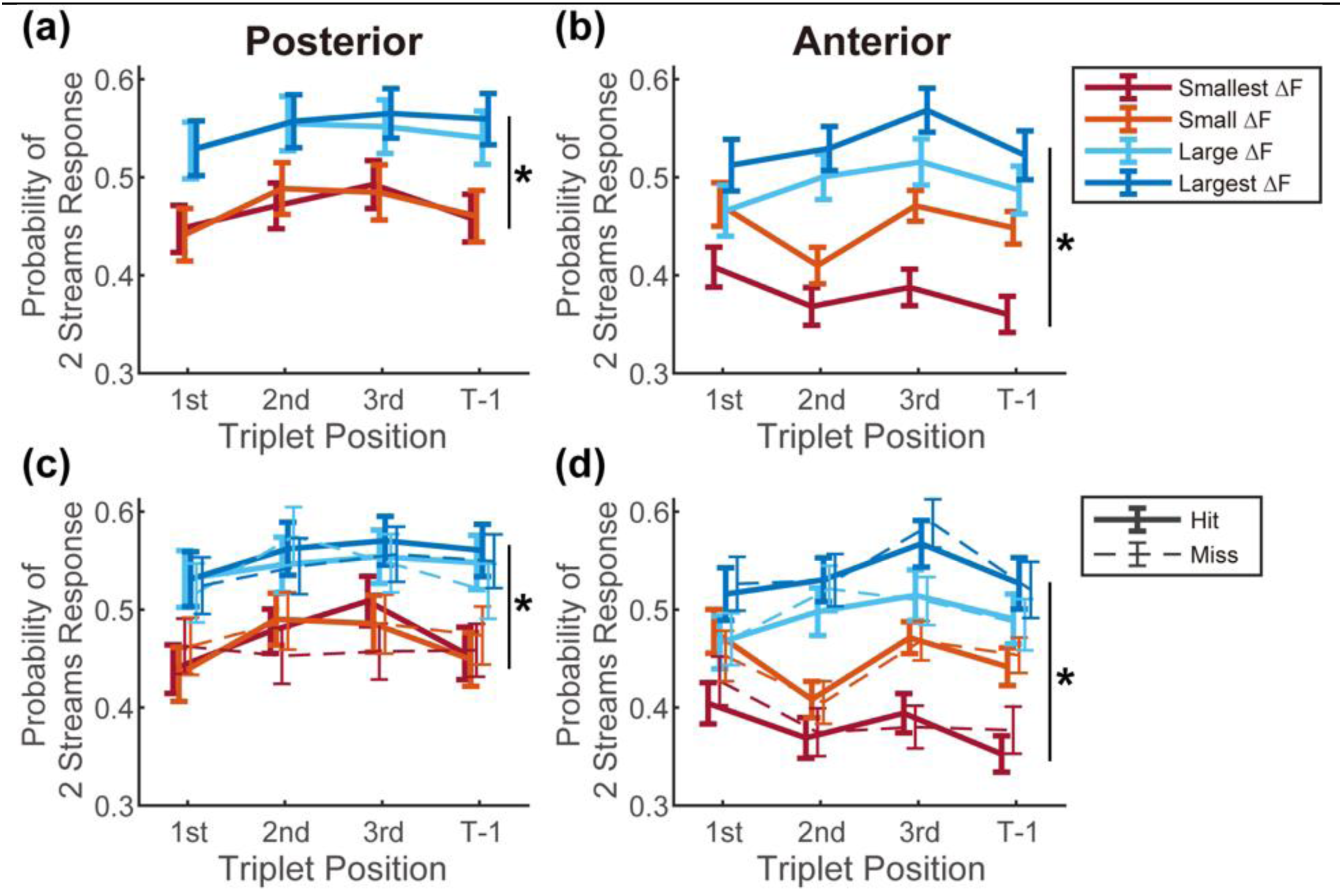
Ideal-observer model can predict “one stream” versus “two stream” choices consistent with behavioral choices. Each panel plots the fraction of trials in which an ideal-observer model classified neural activity “two streams”, as a function of time. Data are aligned relative to the first three (“1^st^”, “2^nd^”, and “3^rd^”) triplet positions and for the triplet immediately preceding the target (i.e., “T-1”). The probability of “two stream” choices from the model trained on data from the posterior auditory field are shown in **(a)**, whereas the probability of “two stream” choices from the anterior auditory field are shown in **(b)**. Data are plotted as a function of ΔF (see legend) but independent of behavioral choice. Data in **(c)** and **(d)** show the probability of “two stream trials” from data generated on hit (solid line) and miss trials (broken line), across all ΔFs from the posterior and anterior auditory fields, respectively. In each panel, data points indicate the population mean and standard error. The asterisks indicate when the null hypothesis was rejected (p<0.05, H_0_: probability of “two stream” reports is the same across ΔF; Scheirer-Ray-Hare test).

### Choice-related activity in the posterior and anterior auditory fields

Finally, in a direct test of choice-related activity, we conducted another auROC analysis to examine whether zMUA was differentially modulated by hit and miss trials (Fig. 7). We found a significant increase in choice-related activity for all three tone bursts in an L-H-H triplet (p = 3.12 x 10^-18^ for low-frequency tone, p = 1.24 x 10^-25^ for the first high-frequency tone burst, p = 5.98 x 10^-17^ for the second high-frequency tone burst; H_0_: auROC values were the same across triplet position; Scheirer-Ray-Hare test) in both the posterior and anterior auditory fields (Table 3). Early in the streaming sequence, the auROC values were close to 0.5 (i.e., no difference between hit and miss trials). As the trial unfolded, auROC values increased slightly and became greater than 0.5 (Table 4; p<0.05; Wilcoxon signed-rank test, FDR corrected), especially during the target triplet and the one immediately following it. We also found some evidence that choice-related activity (which was elicited by the low-frequency and first high-frequency tone bursts) was greater in the anterior auditory field than in the posterior auditory field (p = 0.007 for low-frequency tone burst, p = 0.018 for the first high-frequency tone burst; H_0_: auROC values were the same in both auditory fields; Scheirer-Ray-Hare test) but not for the second high-frequency tone burst (p = 0.08, Scheirer-Ray-Hare test).

**Figure 7:**
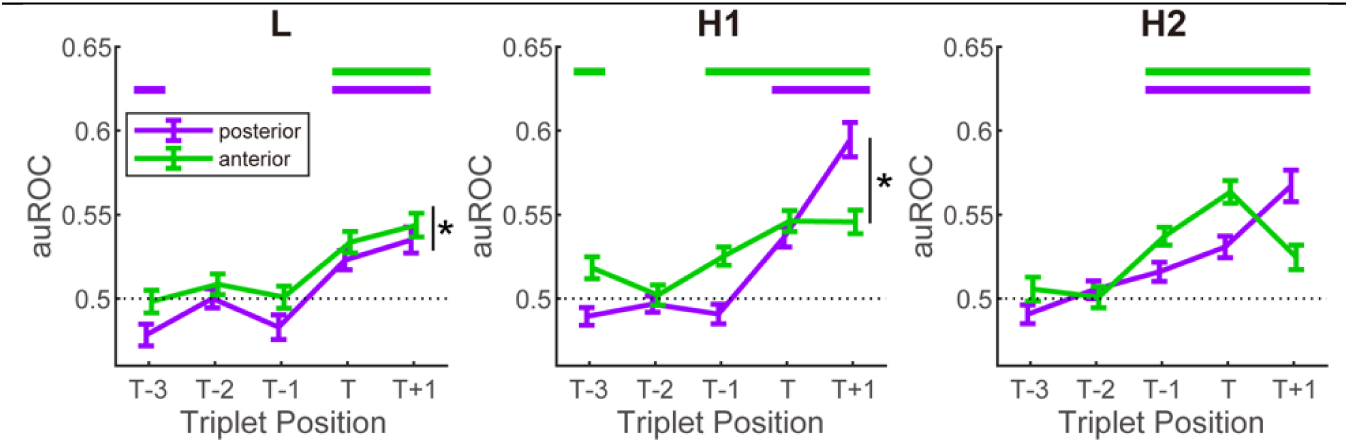
Choice-related modulation of neural activity increases prior to target onset. auROC values plotted as a function of target onset, starting with the three triplets preceding target onset (respectively, “T-3”, “T-2”, and “T-1”), the target triplet (“T”), and the one following it (“T+1”). auROC values were calculated from distributions of zMUA vales on hit trials and on miss trials as a function of ΔF and then averaged together. auROC values are plotted for data from the posterior (purple) and anterior (green) auditory fields and for (**left**) the low-frequency tone burst, (**middle**) the first high-frequency tone burst, and (**right**) the second high-frequency tone burst. Data points indicate the population mean and standard error. The asterisks when the null hypothesis was rejected (p>0.05, H_0_: choice activity was the same in the posterior and anterior auditory fields; Scheirer-Ray-Hare test).

**Table 3;.** Statistical test examining the time course of choice-related zMUA modulation

| Tone identity | Posterior auditory field | Anterior auditory field |
| --- | --- | --- |
| L | **1.06 x 10<sup>-11</sup>** | **6.95 x 10<sup>-7</sup>** |
| H1 | **1.40 x 10<sup>-23</sup>** | **9.59 x 10<sup>-6</sup>** |
| H2 | **7.70 x 10<sup>-12</sup>** | **4.80 x 10<sup>-10</sup>** |

p-values from Kruskal-Wallis test examining whether the auROC values in Fig. 7 change over the stimulus period including T-3, T-2, T-1, T, and T+1 triplet positions. Bold font indicates statistical significance ( $p < 0.05$ ).

**Table 4;.** Statistical test examining MUA choice sensitivity at each triplet position

| Tone identity | T-3 | T-2 | T-1 | T | T+1 |
| --- | --- | --- | --- | --- | --- |
|  | *Posterior auditory field* |  |  |  |  |
| L | **0.022** | 0.86 | 0.23 | **1.02 x 10<sup>-3</sup>** | **6.98 x 10<sup>-6</sup>** |
| H1 | 0.21 | 0.60 | 0.65 | **3.29 x 10<sup>-7</sup>** | **2.58 x 10<sup>-10</sup>** |
| H2 | 0.23 | 0.60 | **0.039** | **3.57 x 10<sup>-5</sup>** | **1.01 x 10<sup>-7</sup>** |
|  | *Anterior auditory field* |  |  |  |  |
| L | 0.97 | 0.16 | 0.95 | **2.46 x 10<sup>-5</sup>** | **1.35 x 10<sup>-6</sup>** |
| H1 | **0.024** | 0.65 | **2.00 x 10<sup>-4</sup>** | **9.33 x 10<sup>-8</sup>** | **1.12 x 10<sup>-6</sup>** |
| H2 | 0.56 | 0.80 | **1.86 x 10<sup>-7</sup>** | **1.60 x 10<sup>-9</sup>** | **4.31 x 10<sup>-3</sup>** |

p-values from Wilcoxon signed-rank test examining whether the auROC values in Fig. 7 are different from 0.5 at each triplet position. The p-values are FDR corrected. Bold font indicates statistical significance ( $p < 0.05$ ).

## Discussion

We examined how MUA in the posterior and anterior auditory fields of the auditory cortex was modulated during a behavioral task that provided an objective measure of auditory streaming. We observed a reduction in the amplitude of MUA in the posterior auditory field (but not the anterior auditory field) as the streaming stimulus unfolded over time (i.e., at progressively later tone-burst triplet positions within a trial). This reduction depended on the frequency difference between the low- and high-frequency tone bursts of the streaming stimulus. Using a threshold-based neural model of stream segregation similar to that developed by Micheyl and colleagues^7^, we found a pattern of “one stream” and “two stream” decisions that were qualitatively consistent with behavioral results in both humans^1,13, 37–39^ and monkeys^10, 25^. Specifically, the model was significantly more likely to categorize neural activity as “two streams” when ΔF was large than when it was small. Contrary to previous results by Micheyl and colleagues, however, we found that the decision to report “two streams” by the model did not significantly improve or “build up” over time. Further, the model categorization was invariant to the neural activity that was elicited by the monkeys’ different choices.

### Streaming behavior in monkeys is generally comparable to humans

In our previous streaming studies, we trained monkeys to report whether two interleaved sequences of tone bursts were heard as “one stream” or “two streams”^10, 25^. As expected from previous work in human listeners^1,13, 37–39^, the proportion of the monkeys’ reports of “two streams” increased as the frequency difference between the two tone-burst sequences increased. However, an alternative interpretation is that the monkeys were not reporting their streaming percept per se but rather their categorical judgement of whether the tone-burst sequences had a large or small frequency difference.

Here, to rule out this alternative interpretation, we modified the task design so that instead of reporting “one stream” or “two streams”, the monkeys objectively reported a deviantly loud target stimulus (Fig. 1a). This modification was critical because it enabled us to dissociate the stimulus dimension that the monkeys needed to detect (sound level) from the dimension that affected streaming (i.e., the frequency difference [ΔF] of the tone-burst sequences). Indeed, Sussman and Steinschneider (2009)^13^ demonstrated a high correlation between target detection and human participants’ subjective reports of stream segregation. Similarly, on average, our monkeys reported the deviant target more reliably as ΔF increased. Together with previous findings^23–25^, our results add further evidence that non-human primates process auditory streams in a manner that is generally comparable to human listeners.

Although qualitatively similar to those in human subjects^13^, the monkeys’ overall hit rates and d’ values were lower (Fig. 1b and 1c). These behavioral differences may be attributed, in part, to differences in task design. For example, in the human study, the stimulus sequence was presented for several minutes, whereas our stimulus duration was considerably shorter (<2 s). Because stream segregation “builds up” over time^40, 41^, the longer-duration listening time might have given the human listeners a behavioral advantage in detecting the deviant target stimulus. Moreover, in the Sussman and Steinschneider study (2009), during those long stimulus durations, ΔF was held constant with multiple presentations of the same deviant target, whereas we trial-by-trial varied ΔF. Thus, it would be informative to investigate whether the monkeys’ task performance would be more comparable to that of humans with a more similar task design. Nonetheless, the present findings support the utility of Sussman and Steinschneider’s behavioral stimulus paradigm in future studies of auditory streaming in animal models.

### Frequency-dependent modulation of MUA during the streaming task

As the tone-burst triplet sequence unfolded over time, we found a general reduction in zMUA and one that was dependent on the frequency difference between the low- and high-frequency tone bursts (Fig. 4). This reduction in neural responsivity was akin to that seen in previous work^7,14^. One obvious difference was that these previous studies reported more time-dependent neural habituation than suggested by our findings.

However, this may be due to differences in the duration of the sequences: our sequences tended to be much shorter (<2 s) than those of earlier studies (10 s), which limited any time-dependent reduction in neural responsivity. Our stimuli were shorter due to the constraints of having behaving monkeys participate in the streaming task, whereas the previous reports were not engaging monkeys in streaming. In other work in our lab^10^, we noted frequency-independent reduction in neural responsivity but not frequency-dependent, due mostly to the fact that we could not resolve neural responses to individual tone bursts. Nonetheless, our findings suggest that some form of adaptation/habituation occurs in the auditory cortex of behaving monkeys while they perform a streaming task (Fig. 4). We extend previous work by demonstrating not only adaptation/habituation in the primary/core auditory cortex (posterior auditory field) but show that it is also in non-core cortices that overlap with the anterior auditory field.

Could this decrease in responsivity be a mechanism by which the auditory system codes “one stream” versus “two streams” percepts (see Fig. 6)? Specifically, when we set a threshold by which an ideal observer could optimally distinguish between the zMUA elicited by the low-frequency tone burst and the zMUA elicited by the high-frequency tone burst, we found that the probability of reports of “two streams” (i.e., only the low-frequency response is above this optimal threshold) increased with increases in ΔF (Fig. 6). The relationship between ΔF, the probability of reports of “two streams”, and time was comparable in both auditory fields. Interestingly, this read-out was not meaningfully modulated by the monkeys’ actual choices (Fig 6c, d; hits versus misses).

In comparison to a similar model, we did not observe an increase of probability of “two-stream” reports as the sequence unfolded^7^. This may be due to several non-exclusive reasons. First, as noted above, the duration of our stimulus sequences was considerably shorter than those of Micheyl et al.; this is relevant because streaming percepts, especially those of “two streams”, “build up” over time^7,42^.

Second, our study and the Micheyl et al. study might have used different criterion and means to establish the threshold. Finally, the monkeys’ behavioral performance might limit our ability to decode the neural responses, especially as a function of behavioral choice. Nonetheless, at least in our data set, because choice does not modulate read-out substantially, it suggests that (at least at our level of analysis) this auditory activity may not correlate with a listener’s perceptual segregation of auditory streams but may simply reflect bottom-up processes.

### Choice-related modulations in core and belt auditory cortex

We found a comparatively weak but significant choice-related modulations of tone-evoked neural activity in both posterior and anterior auditory fields (i.e., hits versus misses; Fig. 7). Choice-related activity has been reported previously in early auditory fields, such as the core auditory cortex^43–46^. However, whereas activity in the belt auditory cortex has been shown to be causally related to a particular frequency-discrimination decision^47^, it is not clear whether activity in hierarchically ‘earlier’ auditory fields, including core auditory cortex, is similarly part of a feedforward process that underlies an ongoing decision. Indeed, our results indicated that the frequency discriminability of MUA was not significantly modulated by behavioral outcomes (Figs. 5 and 6) even though, at the time of target onset, we did observe choice-related activity both in the posterior and anterior auditory fields (Fig. 7). Choice-related activity was more pronounced in the anterior than the posterior auditory fields, a result which fits into a hierarchical model in which early cortical areas represent lower-level sensory attributes of stimuli and later areas convert these representations into a behavioral decision^47, 48^. However, because it was not possible for us to determine the time of the actual perceptual decision, this choice-related activity, especially that observed in the posterior auditory field, may reflect target anticipation or other top-down attentional processes^49, 50^. Further work that compares the time of the actual perceptual decision with changes in neural activity^47, 51^

as well as causal manipulations would be needed to more fully elucidate the contribution of these choice-related modulations to auditorybehavior and the perceptual organization of sound sequences.

## Acknowledgements

This work was supported by the National Institutes of Health (NIDCD). We thank Dr. Jaejin Lee for help with behavioral training.

